# Gibbs Free Energy of Protein-Protein Interactions reflects tumor stage

**DOI:** 10.1101/022491

**Authors:** Edward A. Rietman, Alex Bloemendal, John Platig, Jack A. Tuszynski, Giannoula Lakka Klement

## Abstract

The sequential changes occurring with cancer progression are now being harnessed with therapeutic intent. Yet, there is no understanding of the chemical thermodynamics of proteomic changes associated with cancer progression/cancer stage. This manuscript reveals a strong correlation of a chemical thermodynamic measure (known as Gibbs free energy) of protein-protein interaction networks for several cancer types and 5-year overall survival and stage in patients with cancer. Earlier studies have linked degree entropy of signaling networks to patient survival data, but not with stage. It appears that Gibbs free energy is a more general metric and accounts better for the underlying energetic landscape of protein expression in cells, thus correlating with stage as well as survival.

This is an especially timely finding because of improved ability to obtain and analyze genomic/proteomic information from individual patients. Yet, at least at present, only candidate gene imaging (FISH or immunohistochemistry) can be used for entropy computations. With continually expanding use of genomic information in clinical medicine, there is an ever-increasing need to understand the thermodynamics of protein-protein interaction networks.

## Introduction

Early understanding about protein-protein interaction (PPI) networks suggest that changes in PPI network architecture correlates with stage[1] and survival[2]. Paliouras et al 2011[1] used mass spectrometry on prostate clinical samples to show how changes in the protein-protein interaction network architecture relate to Gleason score and prostate specific antigen (PSA). Similarly, Freije et al.[2] showed that gene expression profiling of gliomas correlated with patient survival. In order to reduce the uncertainty inherent to a PPI network and ameliorate the difficulties with reconciling disparate PPI networks, one can combine PPI networks, transcriptome, stage and survival data. The consolidation of PPI data with expression transcriptome data into a coherent abstract model is not only likely to improve the quality of the information in each of these previously unrelated data types, but also improve the data quality sufficiently to use the information for personalized therapies.

There are several ways of measuring complexity of protein-protein interaction networks. Recent papers [3, 4] describe *topological metrics* of PPI cancer networks that correlate with 5-yr cancer patient survival. Breitkreutz et al (2012)[5] and Takemoto and Kaori (2013)[6] describe a *thermodynamic measure* based on degree distribution. A degree distribution is essentially a Boltzmann [7] distribution, which allows us to consider real-world thermodynamics.

In the present manuscript we describe a thermodynamic measure of molecular PPI networks. After a brief review of how thermodynamics can be applied to cancer biology, we describe how to compute Gibbs free energy for cancer networks and show its correlation with 5-yr survival and cancer stage.

Thermodynamics and entropy in particular, have been applied to biology and especially to cancer dynamics in the past. In one of the iterations thermodynamics can be applied as information entropy[8], but because there is no intelligent observer in nature in general or molecular information in particular, thermodynamic entropy is the more appropriate measure. Demetrius (2013)[9] reviewed the thermodynamics of biology, and described the directionality in evolution and the manner in which populations of different organisms enable growth of larger populations. An earlier work by Schneider and Kay [10] suggested the role of entropy in large-scale ecosystems. There are clearly similarities in the role of entropy in large ecosystems networks and the role of entropy in protein-protein interaction networks. Tseng and Tuszynski (2010)[11] propose that an understanding of the maximum entropy principle can lead to a better understanding of biological systems. Complex ecosystems maximize entropy to much higher degree and much more rapidly than simple ecosystems.

Thus, the maximization of entropy in complex biological systems parallels maximization of entropy in complex protein-protein interaction networks. They cite examples of small-scale entropy of protein folding, which can be directly correlated to tubulin isotypes in different cancer cell lines, and better define entropy of drug-protein targets. Similarly, earlier manuscripts describing classical Logistic, Bertalanffy and Gompertz models of tissue growth, [12] or using entropy production rate to calculate avascular cancer growth [13] support the relevance of using entropy for network analysis.

At the cellular and tissue level one can calculate the entropy of an individual/collection cell(s) from karyotype and draw a similar analogy of the molecular network interactions. A number of groups have introduced the concept: Davies et al. discusses thermodynamic entropy of self-organization of biological cells and organisms[14], Metze et al. use the same ideas to describe pathophysiology of cancer by calculating the entropy observed in microscopic images of tissues[15], and Castro et al. [16] describes the use of information entropy using karyotypic analysis of 14 different epithelial tumor types. Computing Shannon information from the karyotype they found a Spearman Rho correlation (r_s_ > 0.8) with 5-yr survival of cancer patients[16]. PPI networks are being used with increasing frequency for mining information about cancer dynamics, cancer progression and therapy, but there are no meaningful tools to analyze them. Breitkreutz et al (2012) found a correlation of degree-entropy of PPI with 5-yr survival[5], introducing the concept, and the work was further elaborated on by Takemoto and Kaori in 2013[6]. Thus, the concept of mathematically analyzing complexity of networks is not new. As far back as 1955, Rashevsky suggested that the study of topology can be applied to networks, and introduced degree-entropy as a network complexity measure[17]. His broad thinking in this purely theoretical paper discussed entropy from an information theory perspective, but did not suggest a connection to thermodynamics. The extension of information theory to thermodynamics in networks was made by Dehmer and Mowshowitz (2011), in a review of the varied entropy measures in network analysis[18].

More recently attempts are being made to combine protein-protein interaction network data and RNA expression data. The quest to find correlations between the PPI networks/transcription data and survival/prognosis has continued. In 2012 Liu et al.[19] defined a measure called state-transition-based local network entropy (SNE). It is a Shannon information measure that is probabilistically, or conditionally, dependent on the previous state of a local dynamical network – a Markov process. They used RNA expression data at different stages of tumor development, overlayed it on protein-protein interaction (PPI) network data, and showed that SNE change significantly with cancer progression. Others have used Shannon entropy measure to show that gene expression patterns of melanoma and prostate cancers group according to cancer stage[20]. Shannon entropy, unlike degree entropy is not a thermodynamic measure.

We now introduce Gibbs free energy, a thermodynamic measure encompassing both network complexity and cell thermodynamics (as represented by transcriptome), and show that it can be correlated with cancer stage and survival.

## Theoretical Background

The homeostasis of cells is maintained by a complex, dynamic network of interacting molecules ranging in size from a few dozen Daltons to hundreds of thousands of Daltons. Any change in concentration of one or more of these molecular species alters the chemical balance, or in terms of thermodynamics, chemical potential. These changes then percolate through the network affecting the chemical potential of other species. The end result are perturbations in the network manifesting as concentration changes, giving rise to changes in the energetic landscape of the cell. These energetic changes can be described as chemical potential on an energetic landscape.

Mutational events invariably alter the chemical potential of one or more proteins and/or other molecular species within a single cell. Yet, two neighboring cancer cells in the same microenvironment may exhibit a different energetic landscape because the chemical potential is different within the two cells. Naturally, when a bundle of cells is harvested, for example in a biopsy, and the cells are digested to extract RNA for transcription analysis, the transcriptome is essentially an average of that bundle of cells. Since many genes code for proteins, the transcriptome can act as a surrogate for the concentration of the proteins. To support this conjecture, several research groups have described correlations of mRNA with protein concentrations [21, 22]) and found Pearson correlation, R, to range from 0.4 to 0.8, in a large number of experiments across five different species. More recently studies of the human proteome across multiple tissue types included in the relevant transcriptomic analysis, and found an average correlation between transcription signal and mass spectrometry proteomic information to be 83% [23, 24].

Work more related to our own is Huang et al. (2005)[25] who proposed that RNA expression data are surrogate metrics for the *protein state* of cells and represent the concentration of specific numbers of individual proteins exposed to either dimetholsulfoxide or all-trans-retinoic acid. Thus, the authors first introduced the concept of a chemical energy landscape for cells. Following exposure to the chemical perturbation, the gene expression data were collected at different time points, cleaned to remove low expression genes, and a self-organizing map created. A principal component analysis was then used to produce a map showing the energetic (chemical potential) trajectory of the cells. The transcriptome has been shown to correlate with protein concentrations [23, 24], and can be generally correlated to the state of the cell. Certainly there are high-throughput protein concentration techniques [26], but the transcriptome provides a higher number of measurements (probes) identified with gene label and readily mapped to protein-protein interaction networks (e.g. BioGrid.org).

The dynamics of cells are coordinated and controlled by protein-protein interactions, and the complete set (known) protein–protein interactions (PPI) gives rise to a network. The state-of-the-art database of these PPI networks is Biogrid (<u>http://thebiogrid.org</u>), described by Breitkreutz et al. (2002)[27]. It should be stressed that, even though state-of-the-art today, it is not complete, and does not describe the full species-specific PPI networks. There are several reasons for this, and they include the fact that the proteome has not been fully mapped from open-reading frames to genes and proteins. Consequently, calculations of the networks’ properties such as entropy or the Gibbs free energy should be taken as estimates reflecting the present state of knowledge about these networks.

We report the outcomes of merging two types of data, transcriptome and PPI networks, to compute the energetic state of cancer. We show a correlation between the Gibbs free energy and 5-yr patient survival for different cancers. Similarly, we show a correlation with Gibbs free energy and cancer stage for liver cancer and prostate cancer as two illustrative examples. In the following paragraphs we describe the calculation of Gibbs free energy of cells, outline the data sources, and present the results and discussion. We hypothesize that transcriptome information can be combined with existing PPI networks and calibrated using Gibbs free energy thus improving the quality of the information with the ultimate goal of enabling future use of transcriptomic information for targeted therapies in clinic.

Our basic hypothesis is that protein-protein interaction (PPI) networks, with the transcriptome acting as surrogate to protein concentration, can be used to compute an estimate of the Gibbs free energy of a cell, or a tumor given the available data. Gibbs free energy, by providing a measure of network complexity and robustness can, in turn, predict the success or failure of therapeutic interventions. The interaction network characterizes PPIs with no regard for time, i.e. the network is time invariant and does not show any time dynamics. The implicate interactions can represent either primary or secondary bonding. In either case the reaction can be represented as:

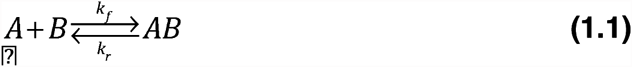

where A and B are two proteins and their interaction product is AB, and *k*_*f*_ and *k*_*r*_ are the forward and reverse reaction rate constants, respectively. When this reaction takes place there is associated with it a bonding free energy (Connors, 1987)[28]. The forward and reverse rate constants are not necessarily equal. From standard physical chemistry we can write the Gibbs free energy of this reaction as: Δ*G* = Δ*H* – *T ΔS*, where the symbols represent the change in Gibbs free energy, G; the change in enthalpy, H, and the change in entropy, S.

Proteins do not interact simultaneously with large numbers of neighbors, as would be implied by the PPI network view of some hub proteins (e.g. p53). Instead the hub protein may be interacting with one or two neighbors at a time forming a complex nanomachine part such as a ribosome. We make the ensemble assumption that many copies of the hub protein may be located in many places in cells and each of the copies may be interacting with a different protein partner. Therefore, we can *assume* an ensemble of the protein of interest, as well as that its interactions with its neighbors are akin to an *ideal gas mixture*.

To help in the understanding of the calculation of Gibbs free energy from the transcriptome and the PPI, we present a simple example shown in Figure 1. Figure 1 shows a small network with individual nodes (proteins) within the network (labeled A, B, C, D, E, and F). For example, D represent a protein connected to E, C, and F by its edges (or links), which represent the interactions between the proteins. Because there is no directionality assigned to the links, the network is said to be an undirected. We compute the Gibbs free energy for protein D below. The network reveals that protein D interacts with proteins C, E, and F, and assuming an ideal mixture of these three proteins, we can assign a nominal chemical potential:

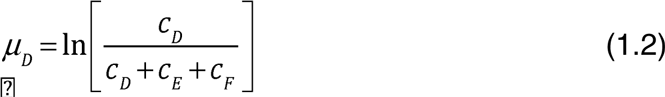

where *c*_*i*_ denotes the concentration of protein *i*. Since Eq. (2) is written as a ratio, we can replace the concentrations with mole fractions, or even normalized expression, to give the same chemical potential. This is known as the entropy of mixing (Maskill, 1985)[29]. The nominal chemical potentials, represented with either concentration or expression, can be used to calculate a nominal Gibbs free energy for not only a single protein with its neighbors, but also for the entire network, for the cell, and the tumor as represented by the transcriptome.

**Figure 1:**
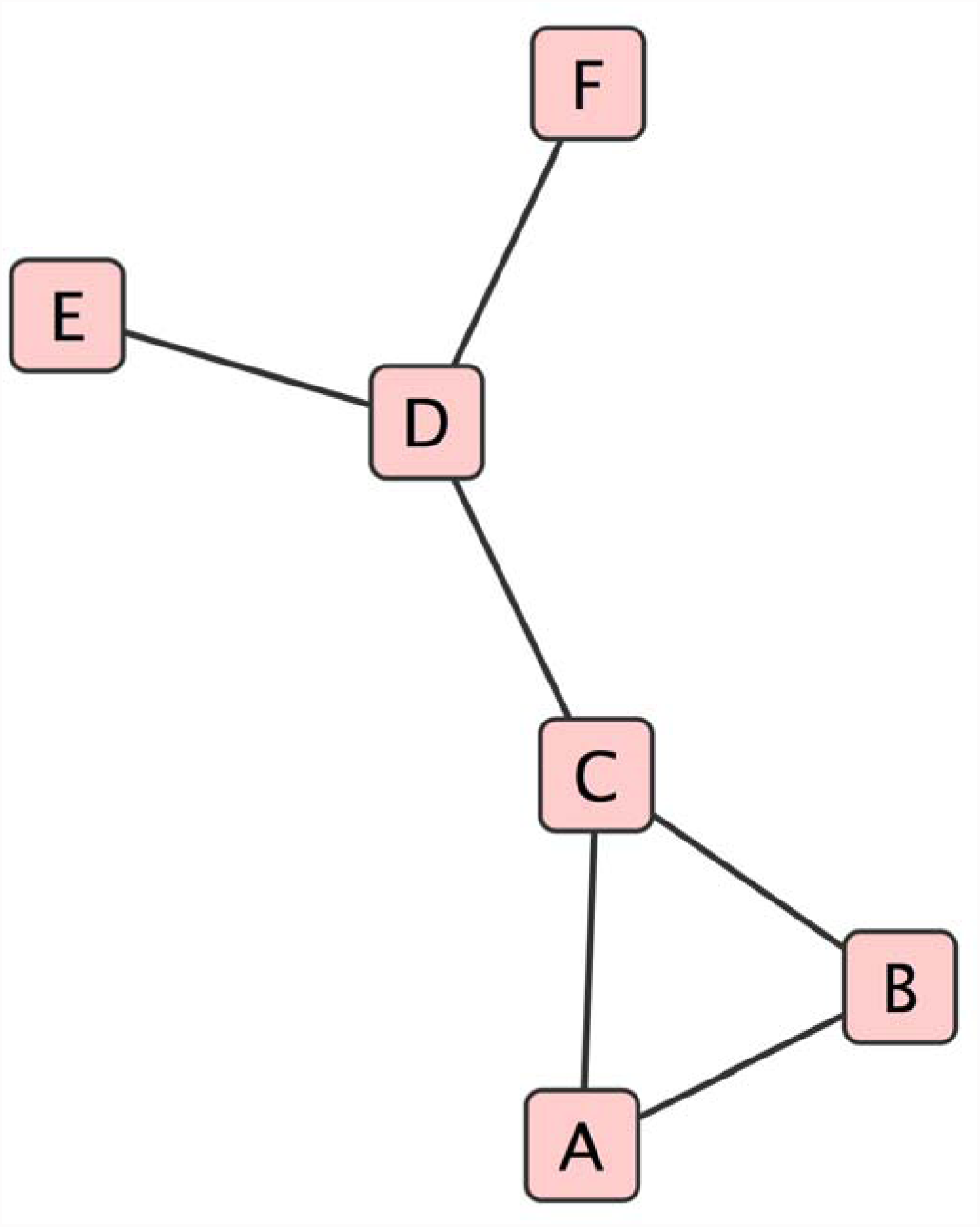
An example of a small protein-protein interaction network created using Cytoscape®. The nodes (A-F) represent individual proteins, the lines, called edges, represent protein-protein interactions. No information about directionality of the interactions is implied. Protein D, for example, represent a protein connected to E, C, and F by its edges (or links). To compute Gibbs free energy for node D in this network, we start with the normalized gene expression data as a surrogate for protein concentration of each node in this network. Gibbs free energy for node D would be: normalized gene expression value divided by the sum of normalized expression of node D+the normalized gene expression values of the neighbors (E,F,C). This quotient becomes the argument for the natural logarithm. The coefficient of the natural logarithm is the normalized expression value for node D. All this is summarized in equation 2.

The chemical potential can be used to compute the Gibbs free energy for node D in the above network as follows:

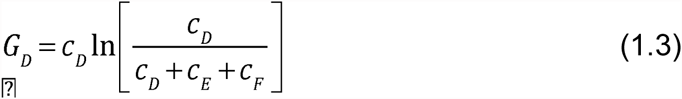

Gibbs free energy scales the expression to thermal energy units, and we can drop the usual convention of including the RT coefficient. Furthermore, because we do not have information on the molar fractions, or molar concentrations, we substitute a normalized, (rescaled) [0,1] RNA transcription value in place of the concentrations.

The general equation for Gibbs free energy can thus be written as:

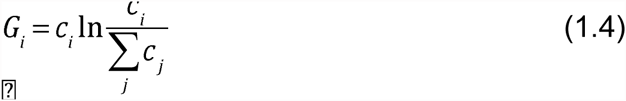

Where the sum is over all neighbors *j* to node *i*, and the sum includes the concentration of node *i*. We can now compute this quasi-Gibbs free energy for the tumor by summing over all the nodes in the network:

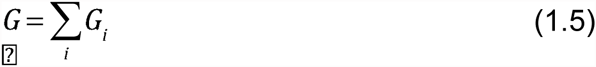

## Data Sources and Methods

Data for several cancers from The Cancer Genome Atlas (TCGA) hosted by the National Institute of Health (<u>http://cancergnome.nih.gov</u>) were collected. The Cancer Genome Atlas is described in TCGA-Research Network, et al., (2013)[30]. More specifically, we collected a set of data that used the Agilent platform G4502A and was pre-collapsed on gene symbols. We collected a total of eleven cancers: KIRC (kidney renal clear cell, TCGA 2013b)[31]; KIRP (kidney renal papillary cell); LGG (low grade glioma); GBM (glioblastoma multiforme, TCGA, 2008); COAD (colon adenocarcinoma, TCGA 2012a); BRCA (breast invasive carcinoma, TCGA 2012c)[32]; LUAD (lung adenocarcinoma); LUSC (lung squamous cell, TCGA 2012b)[33]; UCEC (uterine corpus endometrial, TCGA, 2013a)[34]; OV (ovarian serous cystadenocarcinoma); READ (rectum adenocarcinoma).

We used the human protein-protein interaction network (Homo sapiens, 3.3.99, March, 2013) from BioGrid, which contains 9561 nodes and 43,086 edges. BioGrid (<u>http://thebiogrid.org</u>) [35, 36]. The entire human PPI was loaded into Cytoscape (version 2.8.1[37]). The list of genes obtained from TCGA (full-length expression set was 17,814 genes) for a specific cancer was “selected” using the Cytoscape functions, the “inverse selection” of Cytoscape function applied, and the nodes and their edges were removed. The resulting network, which now included only those genes found in both Biogrid and TCGA, consisted of 7951 nodes and 36,509 edges. This Cytoscape network was unloaded as an adjacency list for processing by custom Python code using Python (2.6.4) with appropriate NetworkX functions.

We used two databases for survival data: The Surveillance Epidemiology and End Results (SEER) National Cancer Institute database, which contains detailed statistical information about the five-year survival rates of patients with cancer, and the National Brain tumor Society database.

For transcription data relevant to prostate and liver carcinoma, we accessed Gene Expression Omnibus (GEO) at <u>http://ncbi.nlm.nih.gov</u>. The data for the liver cancer study (hepatocellular carcinoma) was GSE6764[38], and the prostate study GSE3933 [39] and GSE6099 [40].The GSE3933 and GSE6099 as obtained were log-2 processed, and collapsed to gene IDs. The data was modified to gene cluster text file format (.gct) format and processed with GenePattern® at Broad Institute. The expression data for liver cancer, GSE6764, was in an Affymetrix® format (HG_U133_Plus_2 probe set), and also preprocessed to collapse them into gene IDs. The GSE6764 dataset, the liver data, were not preprocessed by log-2. Consequently the numerical value of the Gibbs energies between those data that were log-2 processed and those data that were not differ and are not comparable. Nonetheless as we show below that preprocessing is not important for scaling between 0 and 1 for concentration.

## Results

Using equations [4,5] we computed the Gibbs free energy for each node in the network as well as the sum of all nodes, i.e. total Gibbs free energy. The analysis is limited to cancers for which transcription data existed in the TCGA database. All of the data sets had used the Agilent® platform, providing a very good gene ID match across all cancers listed in Table 1. The data, which were already log_2_ transformed and collapsed into gene IDs, were averaged across samples for each gene to create a single expression vector representing the entire set for each cancer. To evaluate correlation of Gibbs free energy with cancer stage, we calculated an average expression vector for each stage. Table 1 also shows the number of samples, the types of cancers and the respective survival rates.

**Table 1:**
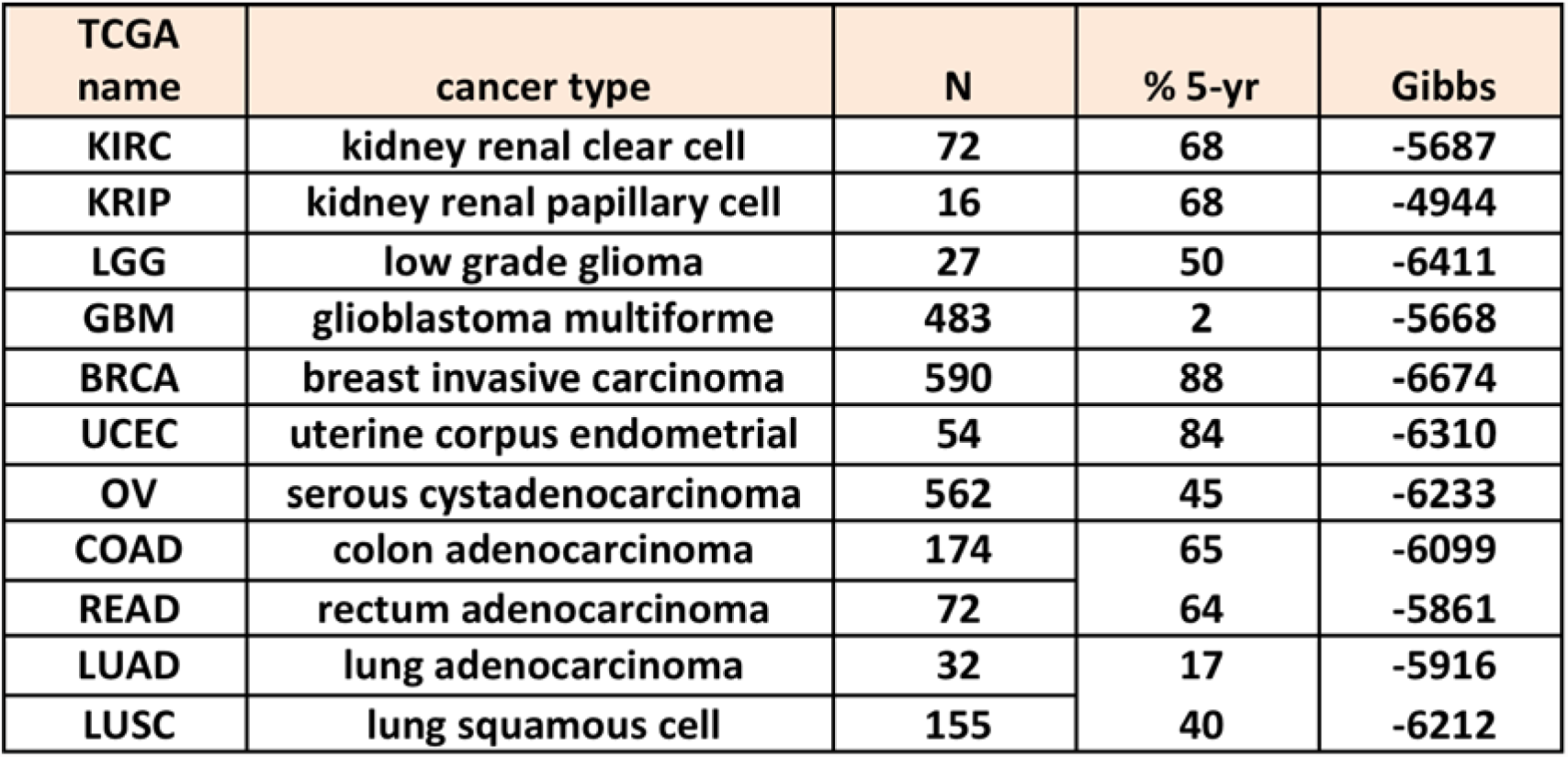
Summary table of the number of subjects in TCGA data sets and respective 5-year survival of individual cancer types from SEER. We collected a total of eleven cancers: KIRC (kidney renal clear cell, TCGA 2013b)[31]; KIRP (kidney renal papillary cell); LGG (low grade glioma); GBM (glioblastoma multiforme, TCGA, 2008); COAD (colon adenocarcinoma, TCGA 2012a); BRCA (breast invasive carcinoma, TCGA 2012c)[32]; LUAD (lung adenocarcinoma); LUSC (lung squamous cell, TCGA 2012b)[33]; UCEC (uterine corpus endometrial, TCGA, 2013a)[34]; OV (ovarian serous cystadenocarcinoma); READ (rectum adenocarcinoma). Gibbs free energy included in this table is the average of the respective N for each individual cancer and was computed using equation 4,5.

| TCGA name | cancer type | N | % 5-yr | Gibbs |
| --- | --- | --- | --- | --- |
| KIRC | kidney renal clear cell | 72 | 68 | -5687 |
| KRIP | kidney renal papillary cell | 16 | 68 | -4944 |
| LGG | low grade glioma | 27 | 50 | -6411 |
| GBM | glioblastoma multiforme | 483 | 2 | -5668 |
| BRCA | breast invasive carcinoma | 590 | 88 | -6674 |
| UCEC | uterine corpus endometrial | 54 | 84 | -6310 |
| OV | serous cystadenocarcinoma | 562 | 45 | -6233 |
| COAD | colon adenocarcinoma | 174 | 65 | -6099 |
| READ | rectum adenocarcinoma | 72 | 64 | -5861 |
| LUAD | lung adenocarcinoma | 32 | 17 | -5916 |
| LUSC | lung squamous cell | 155 | 40 | -6212 |

Before actually overlaying the expression data on the PPI network the average expression vector is rescaled to be in the range [0,1], effectively setting highly up-regulated gene expressions to 1 and highly down-regulated gene expressions to 0. A base assumption was made that previously established correlation that highly up-regulated genes result in a high protein concentration and highly down-regulated genes result in a very low protein concentration [23, 24]. This prevented any negative argument in the natural logarithm of Equation [4], and provided consistency from a chemical physics perspective. The calculated Gibbs values are shown in Table 1.

A plot of Gibbs free energy values versus percent 5-yr survival for these cancers is shown in Figure 2. There are nine cancers shown in the graph (GBM, LUAD, LUSC, READ, COAD, OV, LGG, UCEC, BRCA) with Pearson R correlation of - 0.718, with a p-value 0.0294. KRIC (“Kidney renal clear cell”) and KRIP (“kidney renal papillary cell”) are abnormal tissue growths, which, even though highly proliferative and destructive, are of questionable malignant potential. If one were to include these two abnormal growths (KRIC and KIRP) in the analysis, the correlation would drop to -0.016.

**Figure 2:**
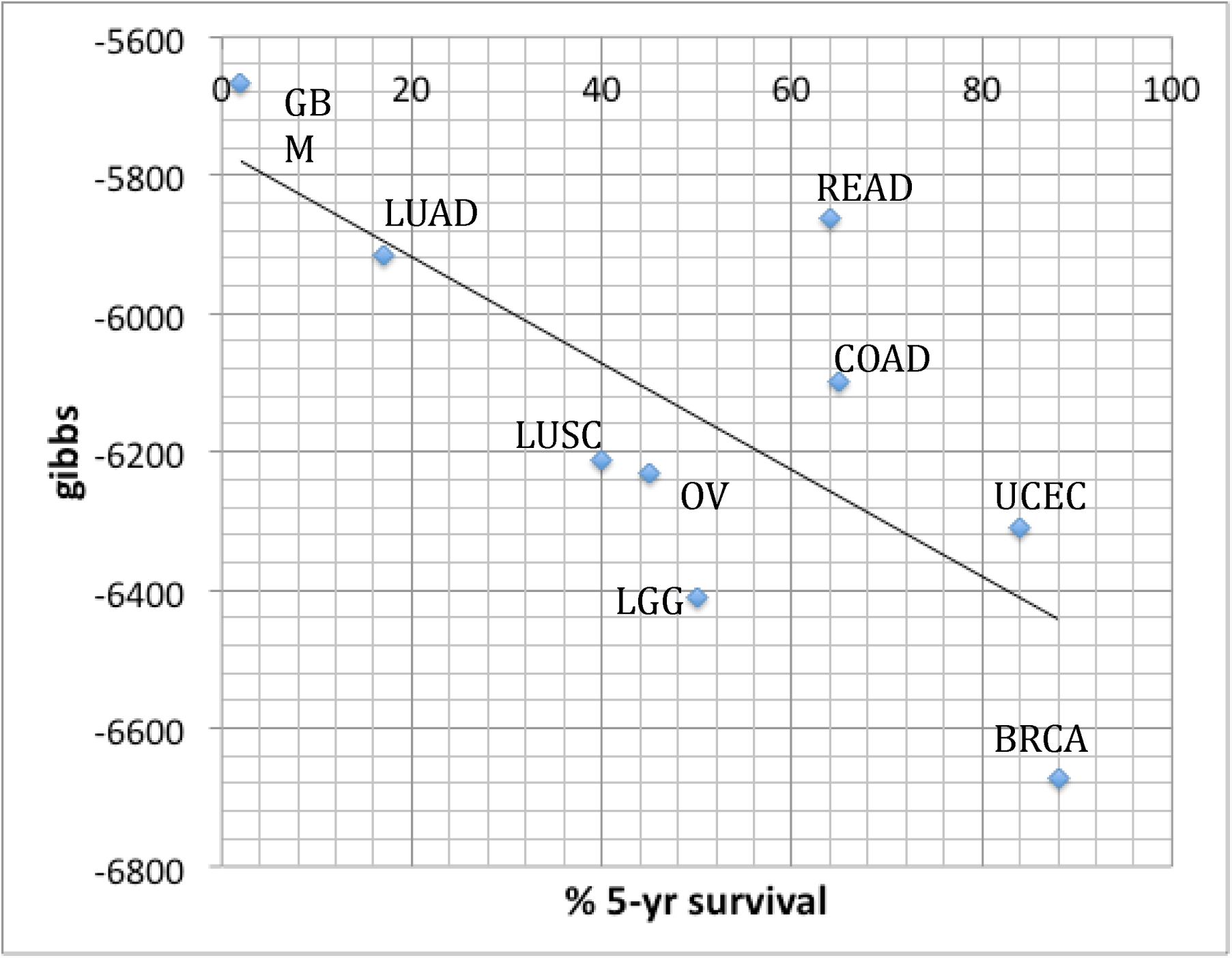
Gibbs free energy and the probability of 5-yr survival. Data from the TCGA gene list were overlaid on BioGrid® in order to merge protein-protein interaction network data with transcription data using Equation 4. As evident, Gibbs free energy can be correlated with 5-year survival with an Pearson R coefficient of -0.718, p-value 0.0294. We have excluded KIRC and KIRP, because the biology of neuroectodermal and epithelial cancers differ from KIRC and KIRP. The inclusion of KIRC and KRIP in the calculation decreased correlation to -0.016.

For comparisons, we compared another measure of the expression data versus survival. We calculated singular values using numpy.lanalg.svd(X) in Python and compared them to survival. The first three singular values versus survival gave R correlations of: -0.070, +0.115, +0.176, respectively (leaving out KIRC, KRIP). These are very poor correlations, and it is reasonable to conclude that Gibbs free energy is more effective in evaluating a real effect on survival or cancer stage, because it is associated with significant changes in energy. An important implication of the correlation between Gibbs free energy and survival/stage is that the higher the Gibbs free energy of a given cancer cell, the more robust it is against external perturbations and the lower the probability of patient survival over a 5 year period. In other words, relative robustness of a cancer cell type can be a prognostic measure of the malignant phenotype of the cancer. This is consistent with other concepts in physics where Gibbs free energy is a measure of stability of a thermodynamic system. Gibbs free energy and entropy are both thermodynamic measures, and because the observations are similar, we can compare the two thermodynamic measures.

As noted in the Introduction, the degree distribution used by Breitkreutz et al. (2012)[5] is essentially a Boltzmann distribution. This allows us to compare entropy with Gibbs free energy. The empirical equation for the linear fit of the Gibbs free energy with survival without kidney cancer is: G = 8.112σ + 5753.9 (Figure 2). Using the data from Breitkreutz et al. [5] we can write the empirical equation for the liner fit of entropy as: *S* = -0.0087σ + 2.2731. Solving both these equation for 5-year survival probability, σ, and equating we get: □ = 7873 - 932□. Note that in order to relate *G* and *S*, we used the absolute value of the Gibbs. This is consistent with the fundamental thermodynamic relationship linking Gibbs free energy and entropy: *G=H-TS*. What remains to be analyzed in the future as more data sets become available is the nature of the proportionality constant playing the role of the absolute temperature, the character of which may be a biological constant of fundamental importance or simply a fitting parameter.

Having shown the correlation of Gibbs free energy with cancer patient survival probability, we turned to examine two specific cancers, stage-by-stage in order to determine whether a relationship exists between the Gibbs free energy and cancer progression. The first cancer analyzed was hepatocellular carcinoma (HCC), one of the more common cancers. We collected GSE6764 data, an Affymetrix data set described by Wurmbach et al. (2007)[38], and processed it using Equations [4,5]. As described by the group contributing the GSE6764 data, three hospitals (Mt. Sinai, New York, USA; Hospital Clinic, Barcelona, Spain; National Cancer Institute, Milan, Italy) were involved in data collection. The results are shown in Figure 3, and define cancer stages as: 0) normal tissue, 1) cirrhotic, 2) low-grade dysplastic, 3) very early hepatocellular carcinoma (HCC), 4) early HCC, 5) advanced HCC, 6) very advanced HCC. The Spearman correlation between these stage-ordinal numbers with respect to Gibbs free energy is R = -1.00 with a p-value of <0.0001 The Kendall tau correlation is - 1.000 and p-value 0.0016.

**Figure 3:**
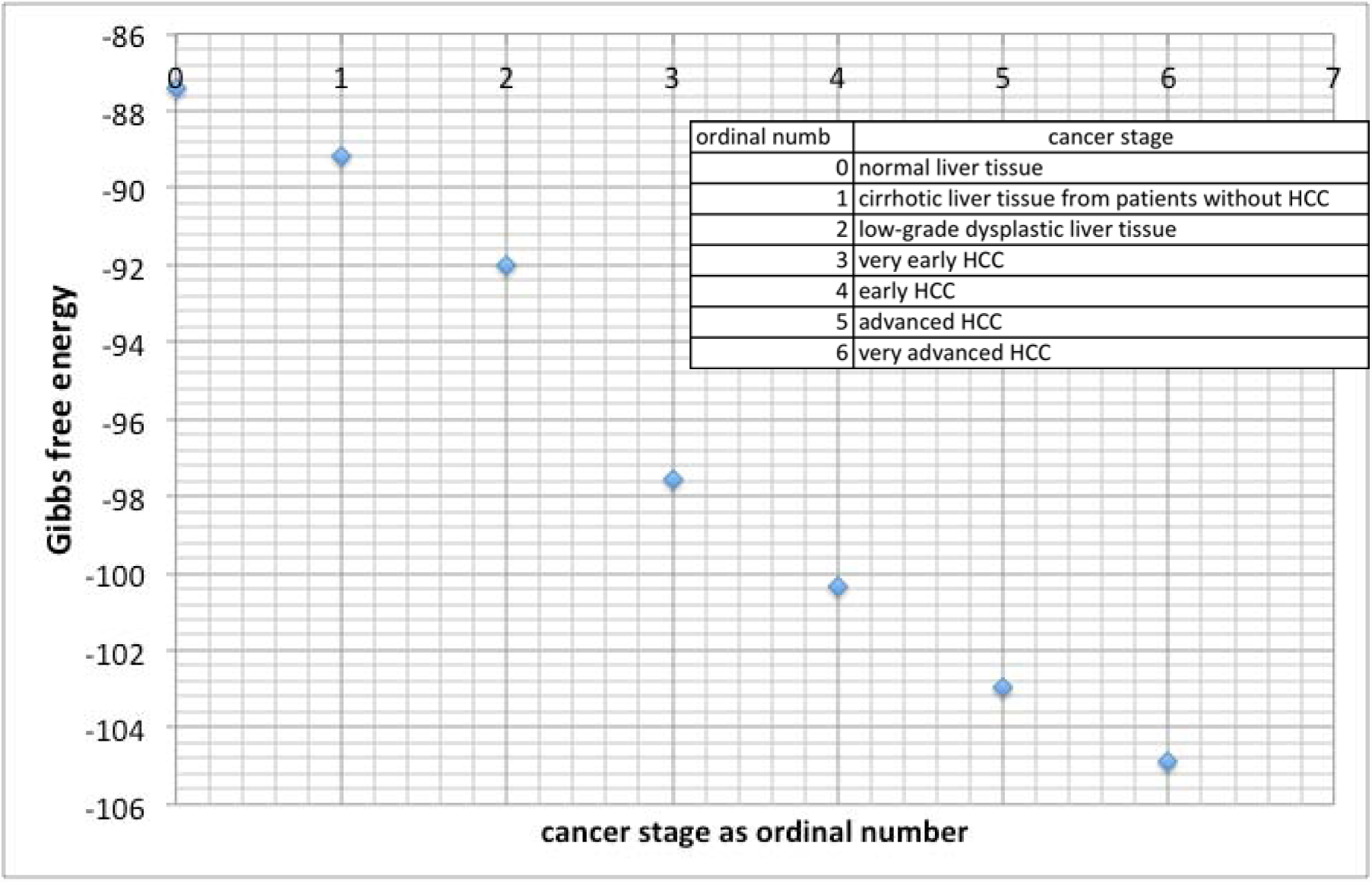
Gibbs free energy correlation with cancer stage for liver cancer. Representing each stage as an ordinal number we present the correlation of: 0) normal tissue, 1) cirrhotic, 2) low-grade dysplastic, 3) very early hepatocellular carcinoma (HCC), 4) early HCC, 5) advanced HCC, 6) very advanced HCC. For this calculation gene expression data from GSE6764 was normalized so as to be in the range of [0,1] and overlaid on a protein-protein interaction network from Biogrid® using equation 5. Unlike the data in Figure 2 or Figure 4, these data were not log-2 preprocessed prior to scaling between 0 and 1. The Spearman correlation of the mean Gibbs free energy for the individual cancer stages is R = -0.99 with a p-value of 0.0001. Kendall’s tau correlation is 1.000, with a p-value of 0.0016.<

The second example was prostate cancers. We collected two completely disparate prostate datasets, one GSE6099 from Lapointe et al. (2004)[39] and another GSE3933 from Tomlins et al. (2007)[40]. The data was compiled, processed as individual transcriptome vectors for computing the Gibbs free energy, and an ordinal integer scale was assigned for each cancer stage. The results are shown in Figure 4, and define cancer stages as: 1) benign prostate hypoplasia (BPH), 2) prostatic intraepithelial neoplasia (PIN), 3) primary tumor, and 4) metastatic disease (MET). The Spearman R correlation is -1.000 with p-value <0.0001. The Kendall tau correlation is -1.000 with p-value 0.0415. Note BPH is essentially age-matched normal prostate tissue for comparison with the diseased tissues. These demonstrate excellent correlation between the thermodynamic measure (Gibbs free energy) and the progression of the neoplastic disease.

**Figure 4:**
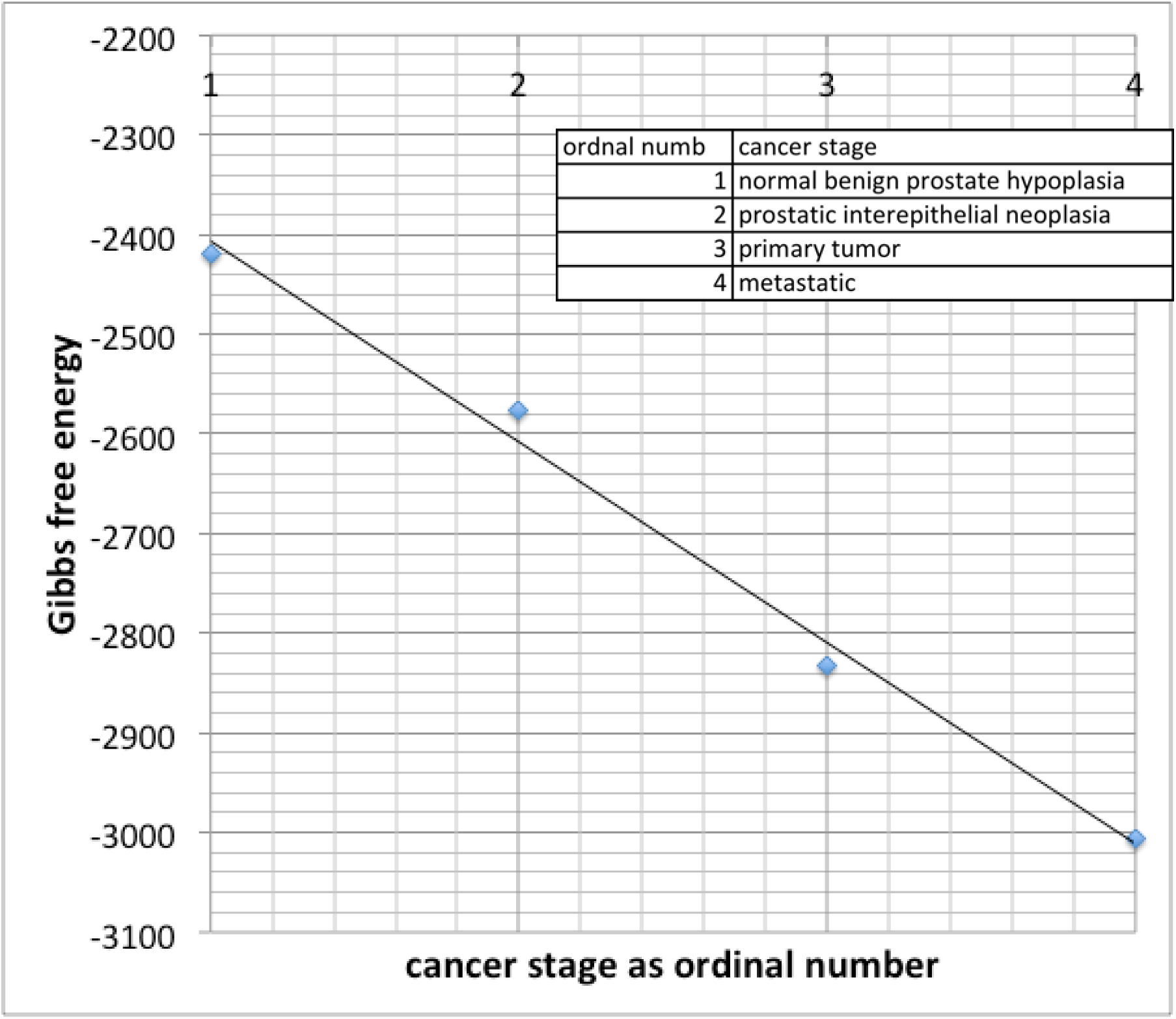
Gibbs free energy vs. cancer stage for prostate cancer. Representing each stage as an ordinal number we present the correlation of normal benign prostate hypoplasia (1), prostatic interepithelial neoplasia (2), primary tumor (3), and metastatic (4). For this calculation gene expression data from GSE3933 and GSE6099 were normalized so as to be in the range of [0,1] and overlaid on Biogrid® protein-protein interaction network using equation 5. The Spearman R correlation is -1.000 with p-value <0.0001. The Kendall tau correlation is -1.000 with p-value 0.0415

## Discussion

As information about cancer related genomic alterations emerge and more and more data becomes available, we can begin to establish the relationships between protein-protein interaction network complexity and cancer progression. We provide Gibbs free energy, a thermodynamic measure encompassing both network complexity and protein concentration (transcriptome), and show that thermodynamics can be correlated with cancer stage and survival. This allows us to potentially differentiate between normal and cancer cells using thermodynamic measures.

We have shown that there is no correlation between the singular values of the expression and survival, and pointed out that the first three singular values (leaving out kidney) versus survival gave R correlations of: -0.070, +0.115, +0.176. This suggests that the expression data is not the most significant component for the analysis and that the PPI network must be playing a significant part. To establish that the network architecture itself does not account for the correlation of Gibbs free energy and survival either, we tested a random network. One can view the mathematical steps in Equations [4,5] as follows:

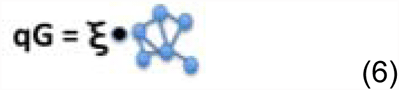

The symbol qG represent a quasi-Gibbs free energy, the symbol ξ represent the expression vector and the little network symbol represents the PPI network. This is analogous to a vector, vector-like product producing a scalar (vector dot product). In these calculations the network architecture is fixed for all expression vectors, for all cancers. To evaluate whether the architecture of the network itself, may play a role, we used random networks, more specifically, random perceptrons (Anderson 1995)[41],and found the dot product for each expression vector with this perceptron network. We computed the indicated dot product, and showed that these random networks did not correlate with survival (R=0.094). Thus, the expression data *and* the PPI network are both needed for a meaningful Gibbs free energy. In effect the PPI network provides a structure to the expression data.

One can use an analogy and view cancer as an invasive species assaulting a complex dynamic ecosystem of the human body – organs and microorganisms all considered. Huang (2011)[42] has argued that the energetic landscape of the epigenome – the epigenetic landscape –inevitably leads to cancer. The molecular network comprising a cell represents a dynamic system on the edge of chaos. Environmental and/or probabilistic fluctuations can push this dynamic system into a trajectory that leads to a stable attractor – cancer – a lower energy state. This concept has been put forward as a general context of cell dynamics by one of the authors of this manuscript[14]. Similarly, Huang et al. (2009)[43] shows how transcriptome data for lung cancer correlates with the various cancer stages, and follows a trajectory of dynamical systems.

A number of other investigators view cancer as an alien species. To name a few, cancer has been viewed as a clonal evolution of cancer cells [44], center around the concept of aneuploidy [45, 46], or creates an analogy that the genome of cancer cells resembles more primitive Metazoa [47]. To maintain their viability cancer cells actually explore a region in attractor space [48] similarly to a strange attractor [49], suggesting many nearby attractors of varying energetic stability. Whatever the forces contributing to cancer evolution may be, they account for observed heterogeneity of cancer cells in the same tumor [50] and provide support to the view that cancer phenotype corresponds to a locally stable Gibbs free energy minimum. Conceptually, a dynamic relationship must exist between stabilizing and destabilizing aneuploidy[48] and the metabolic advantage of tumor cells[51]. This concept of limited attractors in the phase space of cancers is supported by recent research by Hoadley et al. [52] who examined multiplatform data from over 3500 patients and 12 cancer types, and observed that there is only a small subset of mutation types that repeatedly occur in various cancers.

Our work may provide some theoretical support for the recent research reported by Zhang, et al [53] and Suva, et al [54]. The group of Zhang et al [53] describes reprogramming of sarcoma cells in culture. The cells first convert to a pluripotent-like state, and only then differentiate into the appropriate mature connective tissue or red blood cells. Similarly, Suva, et al. [54] describe corresponding reprogramming for the tumor-propagating cells of glioblastoma. We describe cancer as a dynamical system capable of undergoing state changes on an energy landscape. We show this by associating a quantitative measure of the protein-protein interaction network (Gibbs free energy) to the malignancy level of the tumor as a whole (from the transcriptome of tumor biopsy tissues), and show a trajectory from a low-grade tumor to a much higher-grade tumor, as represented by the Gibbs vs. cancer stage plots.. This suggests it may be possible to treat cancer not strictly from a mutation perspective but from an engineering perspective. Rather than simply thinking of inhibiting a specific protein from a mutated gene (or two), it may be possible to treat cancer as a reprogramming of the molecular network with an associated Gibbs free energy landscape. This more holistic perspective considers not just the oncogenes and highly mutated genes but rather the network associated with the relevant proteins and their energetic profile.

## Acknowledgments

EAR was funded by the Newman Lakka Cancer Foundation. JAT acknowledges funding from NSERC, Canadian Breast Cancer Foundation and the Allard Foundation. GLK was funded by NIH NIGMS RO1 GM93050, and Newman Lakka Cancer Foundation. The content is solely the responsibility of the authors and does not necessarily represent the official views of the National Cancer Institute or the National Institutes of Health. We thank Diana White for assistance in compiling data on survival statistics.

## Author Contributions

EAR conceived the idea. JAT and EAR collaborated on the thermodynamics. AB contributed statistical analysis. JP contributed key chemical physics insight. GLK contributed cancer biology expertise. All authors contributed to writing the manuscript.

